# Risk assessment of cancer patients based on HLA-I alleles, neobinders and expression of cytokines

**DOI:** 10.1101/2022.10.15.512339

**Authors:** Anjali Dhall, Sumeet Patiyal, Harpreet Kaur, Gajendra P. S. Raghava

## Abstract

Advancements in cancer immunotherapy have shown significant outcomes in treating various types of cancers. In order to design effective immunotherapy, it is important to understand immune response of a patient based on its genomic profile. We compute prognostic biomarkers from 8346 cancer patients for twenty types of cancer. These prognostic biomarkers has been computed based on i) presence of 352 human leucocyte antigen class-I (HLA-I), ii) 660959 tumor-specific HLA1 neobinders and iii) expression profile of 153 cytokines. It was observed that survival risk of cancer patients depends on presence of certain type of HLA-I alleles; for example LIHC cancer patients with HLA-A*03:01 are on lower risk. Our analysis indicate that neobinders of HLA-I alleles have high correlation with overall survival of certain type of cancer patients. For example HLA-B*07:02 binders have 0.49 correlation with survival of LUSC and −0.77 with KICH cancer patients. It is clear from above analysis that HLA and their binders have major role in survival of cancer patients suffering from different types of cancer. In addition, we compute prognostic biomarkers for 20 types of cancer based on each type of cytokine expression. Higher expression of few cytokines is survival favourable like IL-2 for BLCA cancer patients whereas IL-5R survival unfavourable for KICH cancer patients. In order to facilitate scientific community we developed a web-based platform CancerHLA1 that maintain raw and analyzed data (https://webs.iiitd.edu.in/raghava/cancerhla1/).

## Introduction

Cancer is one of the top causes of mortality worldwide; GLOBOCAN estimates that 19.3 million new cancer cases and 10 million deaths will be reported in 2020 (Sung et al., 2021). Several researchers have worked tirelessly over the last few decades to develop novel cures and treatments to fight against the deadly disease (Pucci et al., 2019). Traditional therapies like chemotherapy, radiation, and surgery are the most commonly used treatments (Arruebo et al., 2011). These radiation-based therapies have negative consequences on the patient’s health and survival (Altun and Sonkaya, 2018;Pucci et al., 2019;Dilalla et al., 2020). To circumvent the drawbacks of conventional medicines, new treatment regimens have been developed, including targeted cancer therapies, adoptive T cell therapy, immune checkpoint inhibitor-based therapies, immunomodulators, interferons and oncolytic viruses (Padma, 2015;Dine et al., 2017;Esfahani et al., 2020;Franzin et al., 2020;Hemminki et al., 2020). Cancer immunotherapy have resulted in considerable outcomes and increase the life duration of many patients suffering from various solid tumours (Amin et al., 2020;Ruiz-Patino et al., 2020).

The central pillars of immunotherapy are immune checkpoint inhibitors and chimeric antigen receptor (CAR) T cells. These therapies are entirely dependent on T-lymphocytes (T cells), which recognise tumor-associated peptides displayed on the tumor cell surface by human leukocyte antigens (HLA) (Waldman et al., 2020). HLAs are the highly complex and polymorphic genes in the human genome, situated on chromosome 6. Class-I HLA alleles interact with the CD8+ T cell receptors to activate T cells which further induce several immune responses to knock off the tumor cells from our system (Buhrman and Slansky, 2013;Engels et al., 2013;Chan et al., 2018;He et al., 2019). The immunotherapies are entirely based on the T cells, which identify tumor-associated peptides presented by human leukocytes antigens (HLA) on the infected cell surface. Recently, scientists majorly focused on HLA-dependent therapies, including CD8+ T cell therapy, tumor-infiltrating lymphocytes (TILs) therapy, and TCR-engineered T cells (TCR-Ts) and neoantigen-based therapy (Sun et al., 2021;Yarmarkovich et al., 2021) to treat cancer patients. HLA-dependent treatments are more effective and efficient than traditional chemotherapies. HLA-peptide binding is critical for determining cancer immunogenicity.

In the past number of repositories such as TCIA (Charoentong et al., 2017), TSNAdb (Wu et al., 2018), NEPdb (Xia et al., 2021), dbpepNeo (Tan et al., 2020), Ovirusdb (Lathwal et al., 2020), CancerTope (Gupta et al., 2016) have been developed for designing immunotherapy. However, these databases lack with information of impact of class-I HLA-alleles, neobinders and cytokines on the survival of cancer patients. To stratify patient-specific therapy, HLA-typing, neoantigens, and binding affinity must be identified. It is now possible to detect patient-specific HLA-alleles with the help of latest technologies and the availability of sequencing data. To construct patient-specific therapy, genetic information such as HLA-alleles, neoantigens, HLA-peptide binding affinity, and immune response need to be considered. In the piolet study, we have developed a resource which provide patient-specific information from public repositories such as (TCGA and TCIA) and analysed patient survival based on HLA-alleles, as well as the correlation of the amount of neobinders specific to HLA-alleles with overall survival in various cancer types. Furthermore, we used correlational analysis to better understand the role of chemokines, cytokines, and their receptors in cancer patient prognosis. We combine all of the aforementioned analysis of 20 types of cancers onto a single user-friendly resource named as “CancerHLA-I” (https://webs.iiitd.edu.in/raghava/cancerhla1/).

## Material and Methods

### Overall study design

The complete architecture of the study is depicted in Figure 1

**Figure 1:**
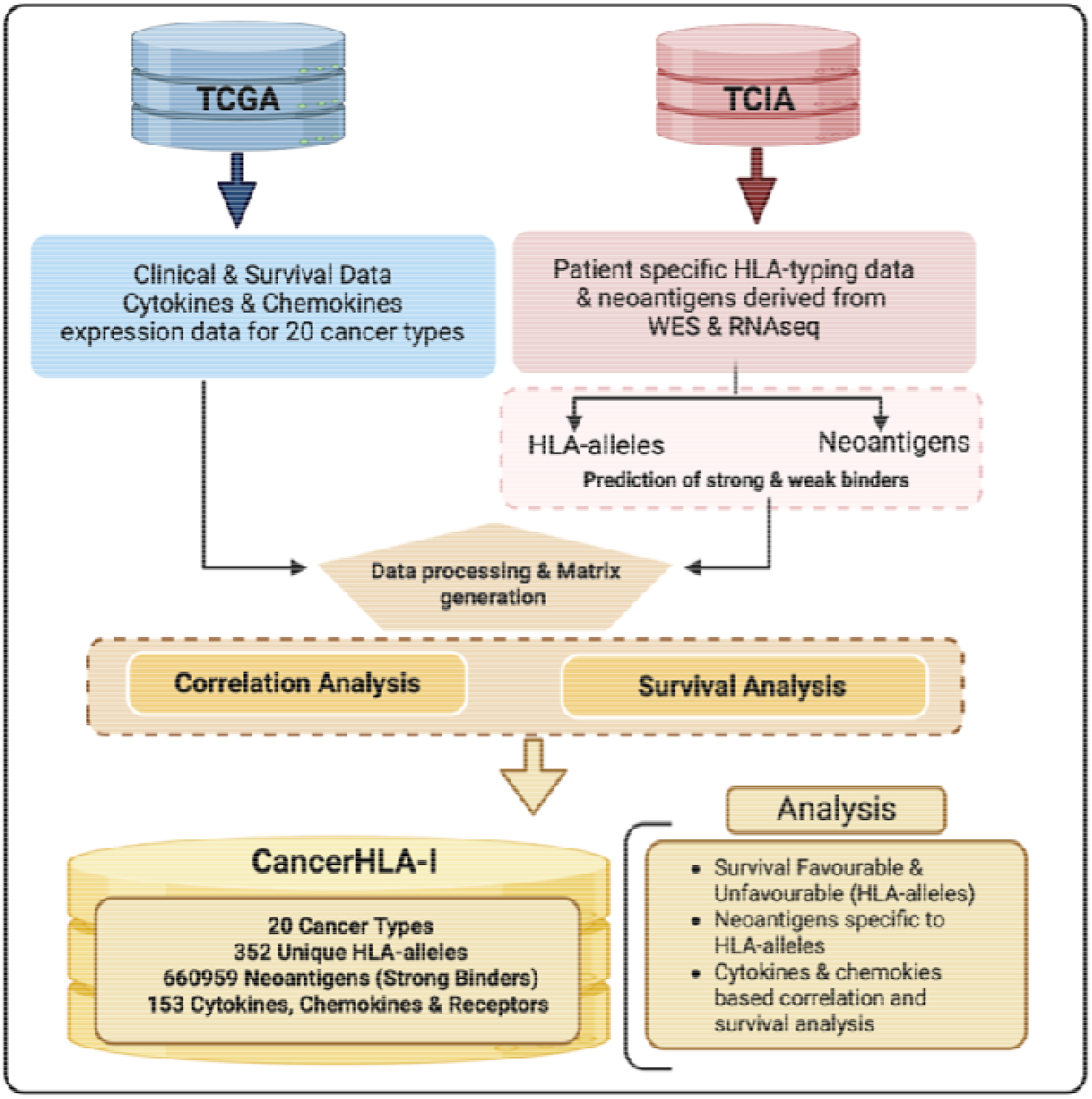
Complete architecture of the study including dataset collection, analysis and website development.

### Collection and pre-processing of datasets

At first, we gathered the genomic and clinical data for 8346 patients with 20 different types of cancer (BLCA, BRCA, CESC, CRC, GBM, HNSC, KICH, KIRC, KIRP, LIHC, LUAD, LUSC, OV, PAAD, PRAD, READ, SKCM, STAD, THCA, and UCEC) from The Cancer Genome Atlas (TCGA) (Tomczak et al., 2015) and The Cancer Immunome Atlas TCIA (Charoentong et al., 2017) repositories. We obtain the control excess dataset patient-specific HLA-typing data and neoantigens data for 20 types of cancers available in TCIA [with the approval of dbGap (Project No. 17674)]. Furthermore, RNA-seq expression data of cytokines, chemokines, and their receptors was obtained using TCGA assembler 2.0. The expression profiles were then transformed into log2 values after addition of 1.0 as a constant number to each of expression value. The survival information covers vital status and overall survival time (OS).

### Mean-overall and univariate survival analysis

For each cancer type, we first built a binary matrix based on the presence or absence of HLA-alleles. Each column represents HLA-alleles, and each row represents samples/patients. We used individual’s survival data to calculate mean overall survival (MOS) based on the presence or absence of an HLA-allele. After that, we computed the difference in MOS (depending on presence/absence). Cox proportional hazard (Cox-PH) regression models were used in the current study to identify HLA-alleles linked with cancer patient survival. The “survival” package in R (V.3.5.1) was used for univariate analysis. The cox regression coefficient greater than 0 indicates that the presence of an HLA-allele which affects survival (unfavourable), whereas less than 0 indicates that the presence of alleles increases survival (favourable). For each allele, we calculated the Hazard Ratio (HR) and 95% CI (Confidence Interval). HR >1 denotes high-risk HLA-alleles, while HR <1 depicts low-risk alleles; however, HR =1 has no effect on survival. Furthermore, the log-rank test and p-value were conducted to determine the significant distribution of low-risk and high-risk patients. To calculate the predictive performance of models, we used the Concordance index (C). In this study univariate survival analysis performed based on HLA-alleles, cytokines, and chemokines genes for 20 each cancer type.

### Correlation analysis

We extracted the strong binding neoantigens/epitopes corresponding to each HLA-allele for each cancer type using the MHCflurry 2.0 software (O’Donnell et al., 2020). We classify neoepitopes as strong or weak binders using the MHCflurry software’s binding affinity (BA) percentile, where neoantigens with BA<2 are considered strong binders and BA>2 taken as weak binders. Following that, we built a count matrix with the number of strong binders matching to each HLA-allele and cancer type. We calculate the correlation coefficient between overall survival and the number of strong binders for each HLA-allele in order to understand the impact of number of binders on the survival using Pearson correlation test. The correlation coefficient and p-value (<0.05) demonstrate the significance of the number of HLA-binding neoepitopes on cancer patient survival. Moreover, we conducted correlation analysis using the expression values of cytokines, chemokines genes for each cancer type. We used survival data and the expression of 153 cytokines, chemokines, and their receptors.

### Database implementation

The web interface of CancerHLA-I (https://webs.iiitd.edu.in/raghava/cancerhla1/) was developed using MySQL and hosted on linux based apache server. In order to create the interactive user interface, we developed responsive framework made up of HTML, CSS, and JavaScript. To improve the data view, the user interface is responsive, which means that the web interface recognises the user device and modifies its structure and shape in accordance with the device resolution. With the help of this functionality, the interface is adaptable to a wide range of devices and browsers with various screen resolutions. The web-site can be searched using variety of devices (smartphones or tablets) and browsers (Google Chrome, Mozilla Firefox, and Safari).

## Results

### Statistical analysis of data of CancerHLA-I

CancerHLA-I incorporates data on cancer associated HLA-alleles, neoantigens, cytokines, chemokine, and their relationship with the overall survival of 20 cancer types. The genomic and clinical information of 8346 cancer patients were downloaded from TCGA and TCIA repositories and processed to build this resource. In Figure 2, we have provided the description of 20 types of cancer, with number of samples, HLA-alleles, total number neoantigens, strong and weak binders. As shown in Figure 2B, HLA-B acquire maximum number of alleles i.e., 185 followed by HLA-A (97 alleles) and HLA-C (70 alleles). We observed that in the case of Uterine Corpus Endometrial Carcinoma (UCEC) highest number (i.e., 192 HLA-alleles) are reported, whereas Kidney Chromophobe (KICH) reported the lowest number i.e., 86 HLA-alleles (See Figure 2C). In addition, we reported top-20 most frequent HLA-alleles whose frequency distribution is maximum among cancer patients (See Figure 2D). Moreover, we have obtained more than 90,000 strong binder i.e., neobinders corresponding to Skin Cutaneous Melanoma (SKCM) and Uterine Corpus Endometrial Carcinoma (UCEC) cancers. However, we have less than 5000 neobinders for Kidney chromophobe (KICH) and Thyroid Carcinoma (THCA) cancer types (See Figure 2E). The complete data used in this analysis is stored in big MySQL table that can be searched and browsed by various categories defined in the database such as cancer type, HLA-allele, neoantigen, HR, p-value, cytokine/chemokine, correlation coefficient etc.

**Figure 2:**
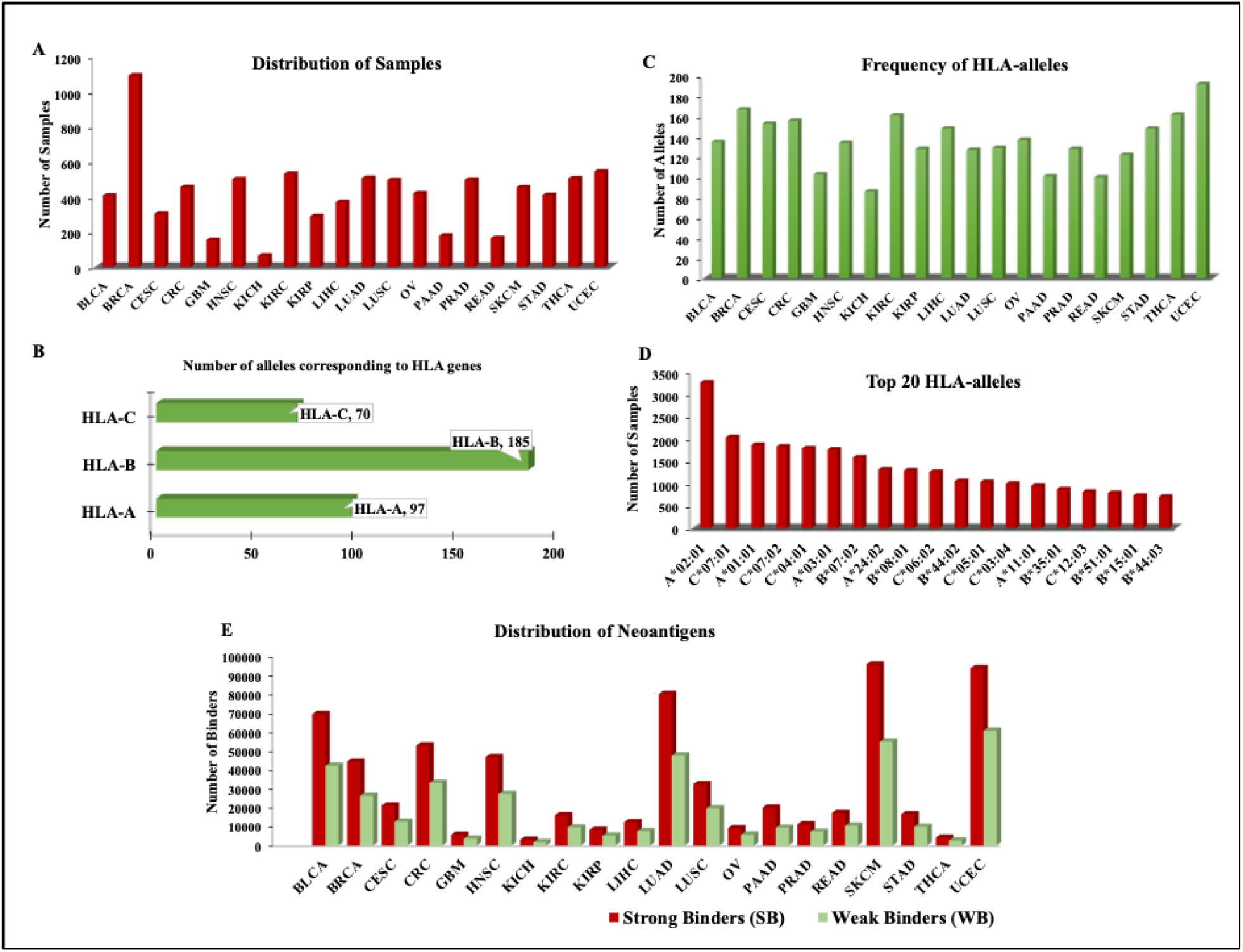
Complete distribution of dataset in 20 cancer types with number of samples, frequency of alleles, HLA-alleles corresponding to class-I genes, most frequent HLA-alleles, and distribution of neoantigens.

Of note, we have generated the UpSet plot to understand the distribution of HLA-alleles in different cancer types. These overlapping and exclusive HLA-alleles among each cancer type (See Figure 3) are the ones that can serve the basis for explaining the molecular heterogeneity and similarity among the different cancer types. It also provides the potential insights into the progression of disease.

**Figure 3:**
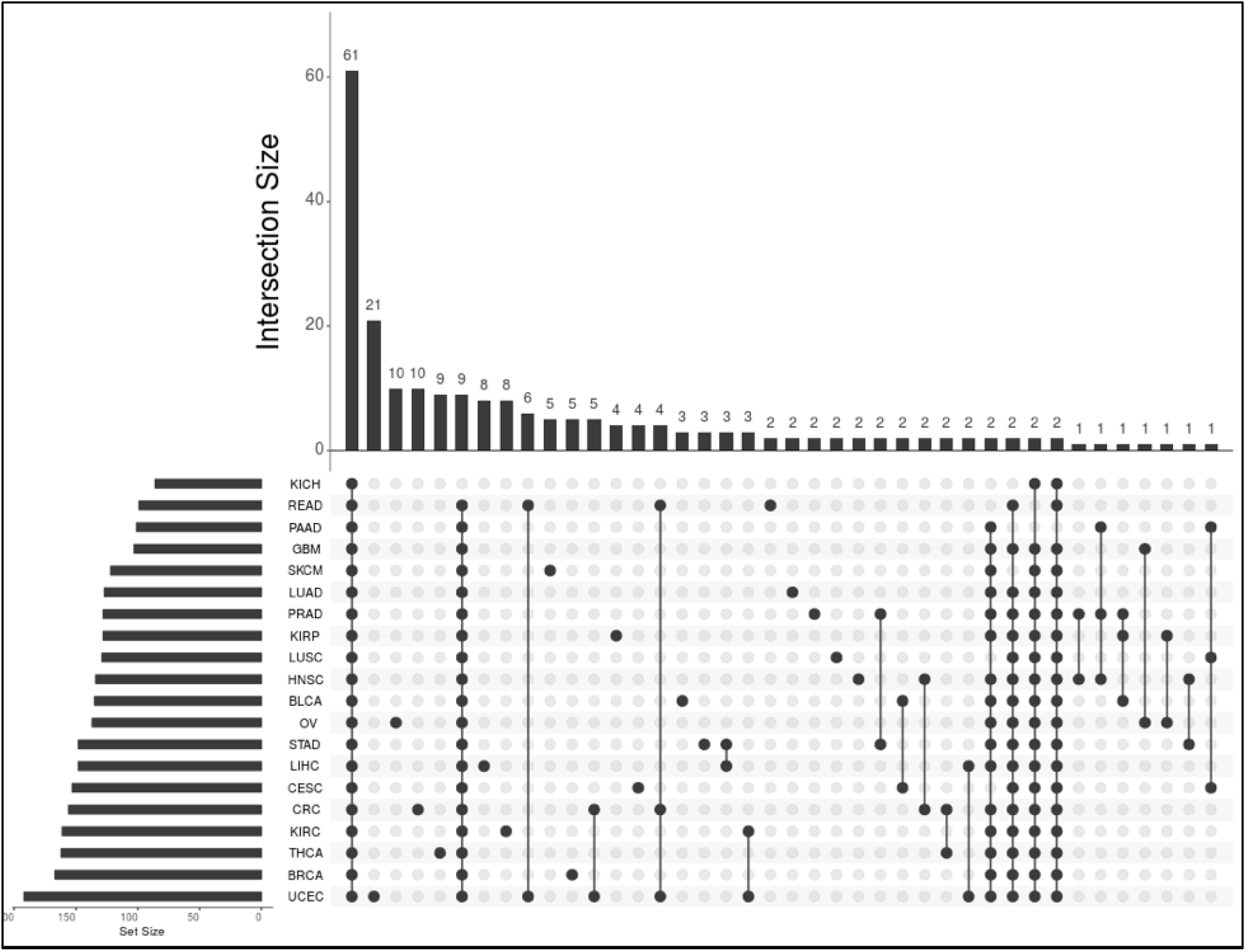
UpSet plot of top 40 interactions including both common and unique HLA-alleles among each cancer type.

### HLA-based prognostic biomarkers

At first, we have combined all the cancer types to performed survival analysis in order to check the overall impact of presence/absence of HLA alleles on the survival of the cancer patients. As shown in the Supplementary Table S1, we have not observed any significant trend by combining all the cancer types due to tumor heterogeneity. So we have generated binary matrix corresponding to each cancer type based on the presence and absence of HLA-alleles. With the utility of survival package we have performed survival analysis computed to identify the high risk and low risk HLA-alleles corresponding to each cancer type. In Table 1, we have reported only those whose HLA-alleles whose presence influences the survival of more than one type of cancer patients. For instance, presence of HLA-A*02:01 allele reduces the survival of kidney cancer (HR=5.46 with p-value=0.03) and skin cancer patients (HR=1.36 and p-value=0.02). We observed that presence of HLA-A*02:01, HLA-A*68:01, HLA-B*52:01 and HLA-C*03:02 is significantly associated with the poor prognosis (with HR>2 and p-value<0.05) in different cancer patients, as shown in Table 1. Moreover, the presence of certain HLA-alleles significantly improves the survival rate of cancer patients for instance, HLA-C*14:02, HLA-C*12:03, HLA-A*03:01 significantly improve the survival rate and act as good prognostic markers in Bladder urothelial carcinoma (BLCA), Stomach Adenocarcinoma (STAD), Liver Hepatocellular Carcinoma (LIHC) and Glioblastoma Multiforme (GBM) (See Table 1).

**Table 1:** Results of univariate survival analysis based on the presence/absence of HLA-alleles in different type of cancers.

| Cancer | HLA-allele | Allele Present<br>(No. of Patients) | Allele Absent<br>(No. of Patients) | Hazard Ratio<br>(95%CI) | p-value | Concordance<br>Index |
| --- | --- | --- | --- | --- | --- | --- |
| KICH | HLA-A*02:01 | 26 | 39 | 5.462 (1.135-26.299) | 0.034 | 0.72 |
| SKCM |  | 203 | 251 | 1.36 (1.038-1.781) | 0.025 | 0.528 |
| CRC | HLA-A*68:01 | 24 | 431 | 2.466 (1.237-4.918) | 0.01 | 0.532 |
| OV |  | 21 | 399 | 1.936 (1.101-3.402) | 0.022 | 0.51 |
| GBM | HLA-B*44:03 | 16 | 138 | 1.805 (1.025-3.181) | 0.041 | 0.528 |
| LIHC |  | 46 | 324 | 1.665 (1.041-2.663) | 0.033 | 0.529 |
| OV |  | 31 | 389 | 1.583 (1.019-2.459) | 0.041 | 0.512 |
| THCA | HLA-B*52:01 | 25 | 480 | 4.057 (1.155-14.254) | 0.029 | 0.62 |
| SKCM |  | 16 | 438 | 2.183 (1.154-4.132) | 0.016 | 0.515 |
| HNSC |  | 18 | 483 | 2.113 (1.149-3.886) | 0.016 | 0.514 |
| LIHC | HLA-C*03:02 | 16 | 354 | 2.268 (1.104-4.658) | 0.026 | 0.517 |
| BRCA |  | 96 | 997 | 1.659 (1.025-2.685) | 0.039 | 0.52 |
| LIHC |  | 43 | 327 | 1.651 (1.034-2.636) | 0.036 | 0.533 |
| LUAD | HLA-C*07:01 | 140 | 367 | 1.366 (0.998-1.869) | 0.051 | 0.538 |
| LUSC |  | 139 | 356 | 1.365 (1.018-1.829) | 0.037 | 0.526 |
| **LUAD** | **HLA-C\*12:03** | 51 | 456 | 1.668 (1.075-2.587) | 0.022 | 0.518 |
| **BRCA** |  | 113 | 980 | 1.655 (1.042-2.627) | 0.033 | 0.543 |
| **GBM** |  | 22 | 132 | 0.527 (0.29-0.936) | 0.029 | 0.532 |
| **KIRP** | **HLA-A\*03:01** | 74 | 215 | 1.884 (1.007-3.526) | 0.044 | 0.531 |
| **LIHC** |  | 72 | 298 | 0.635 (0.397-1.015) | 0.046 | 0.526 |
| **BLCA** | **HLA-C\*14:02** | 15 | 392 | 0.140 (0.02-1.001) | 0.048 | 0.517 |
| **STAD** |  | 20 | 390 | 0.315 (0.1-0.988) | 0.048 | 0.516 |

### HLA-I neobinders based correlation analysis

In this study, we have used MHCflurry 2.0 (O’Donnell et al., 2020) software for the prediction of neoepitopes having strong binding potential with the class-I HLA alleles. We have identified strong HLA-specific neobinders for each cancer type. In order to understand the correlation or impact of number of neobinders with the survival of cancer patients, we performed Pearson correlation analysis. Figure 4, shows the correlation values for the nine HLA-alleles (HLA-A*02:01, HLA-A*03:01, HLA-C*07:01, HLA-C*07:02, HLA-A*01:01, HLA-B*07:02, HLA-B*08:01, HLA-A*24:02, and HLA-B*44:02) present in most of the samples and all cancer types. At first, we have computed the correlation between the neobinders irrespective of cancer type by combining all the data files. We observed that, the overall impact on combining binders for all the cancer types is very less or negligible (See Figure 4). While, some of the neobinders corresponding to particular HLA-allele have very high positive as well as negative correlation with the specific cancer types. For instance, HLA-B*07:02 neobinders have very high negative correlation (r = −0.77) with the survival of KICH patients, while on the opposite side it shows positive correlation of (r = 0.49) with the LUSC patients. In the case of BLCA, LUSC and OV most of the nine alleles shows positive correlation with the overall survival. However, in the case of KICH some of the alleles shows positive association and some of them shows negative association with the survival. This means binders corresponding to HLA-alleles have favourable and unfavourable impact depends upon the cancer types.

**Figure 4:**
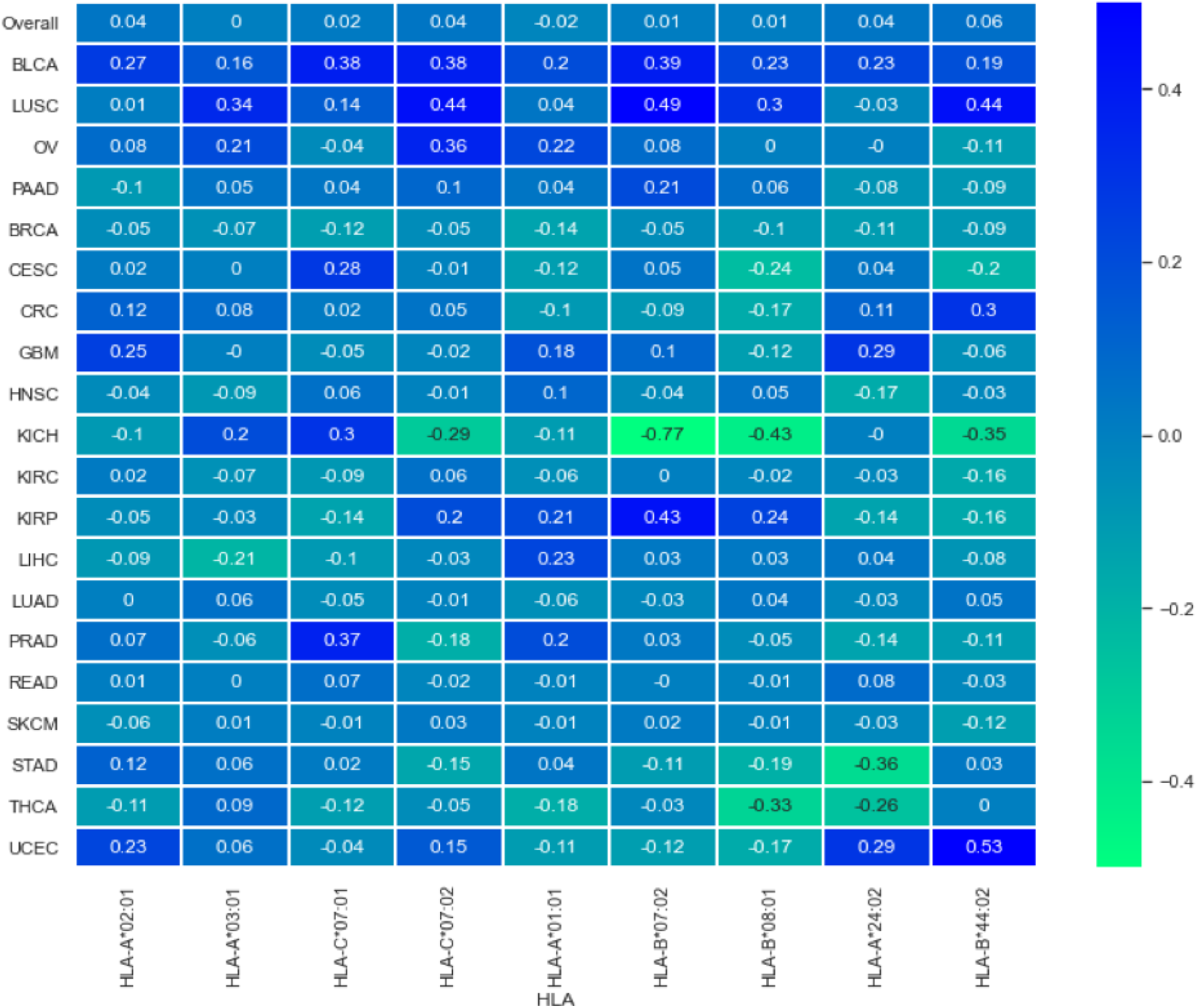
Correlation analysis based on the number of HLA-I neobinders with the overall survival of cancer patients.

### Cytokines based prognostic biomarkers

In order to understand the prognostic role of cytokines and chemokines we have performed univariate survival analysis using their expression profiles. In the Figure 5, we have reported those cytokine and chemokines whose expression significantly impact the survival rate of cancer patients. We observed that high expression of *IL2, IFNB1, IFNA8, IL5* cytokines are having very good impact on the survival of different cancer patients (HR<0.4 and p-value <0.05). Whereas, *IL5RA, TGFBR3, CCR4, TGFB2, IL17A* are highly associated with the poor survival rate in KICH, READ, and GBM patients (HR >4 and p-value<0.05). The complete analysis of all other cytokines and chemokines is available on CancerHLA-I server.

**Figure 5:**
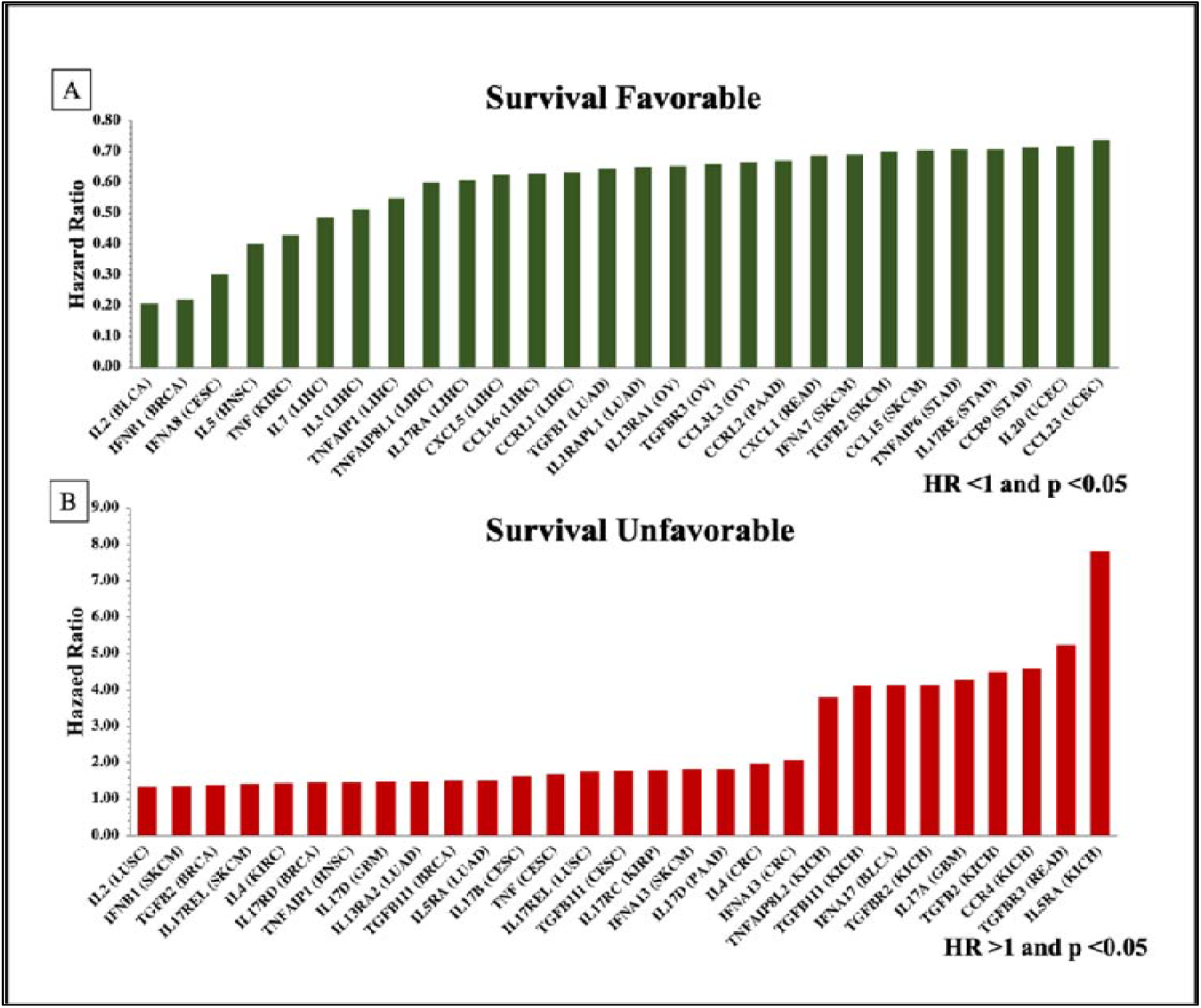
Univariate survival analysis shows survival favourable (HR<1) and unfavourable (HR>1) cytokines, chemokines and their receptors in different cancer types.

Moreover, we have done correlation analysis by considering the gene expression of cytokine, chemokines and their receptors. The heatmap shows (Figure 6) the correlation of overall survival with the expression of some of the cytokines and chemokines in 20 cancer types. The darker blue colour shows the positive correlation, whereas light yellow colour depicts the negative correlation. We observed that cytokine IFNG have very high and significant positive correlation with the survival rate of GBM patients, higher expression of IL9 cytokine is associated with positive correlation in BLCA and OV cancer patients. Whereas, cytokine *IL2, TNFA1P1*, and *TNF* are associated with the negative correlations with the survival of KICH patients. In case of chemokines, we observed that *CCL1* (CRC, KIRC and READ), CCL20 (BLCA, READ, and PAAD), *CCL27* (GBM, KICH and THCA) have positive correlation with the overall survival. However, the over expression of *CCL18, CCL28, CCL4, CCL5* associated with negative correlation with most of the cancer types (See Figure 6).

**Figure 6:**
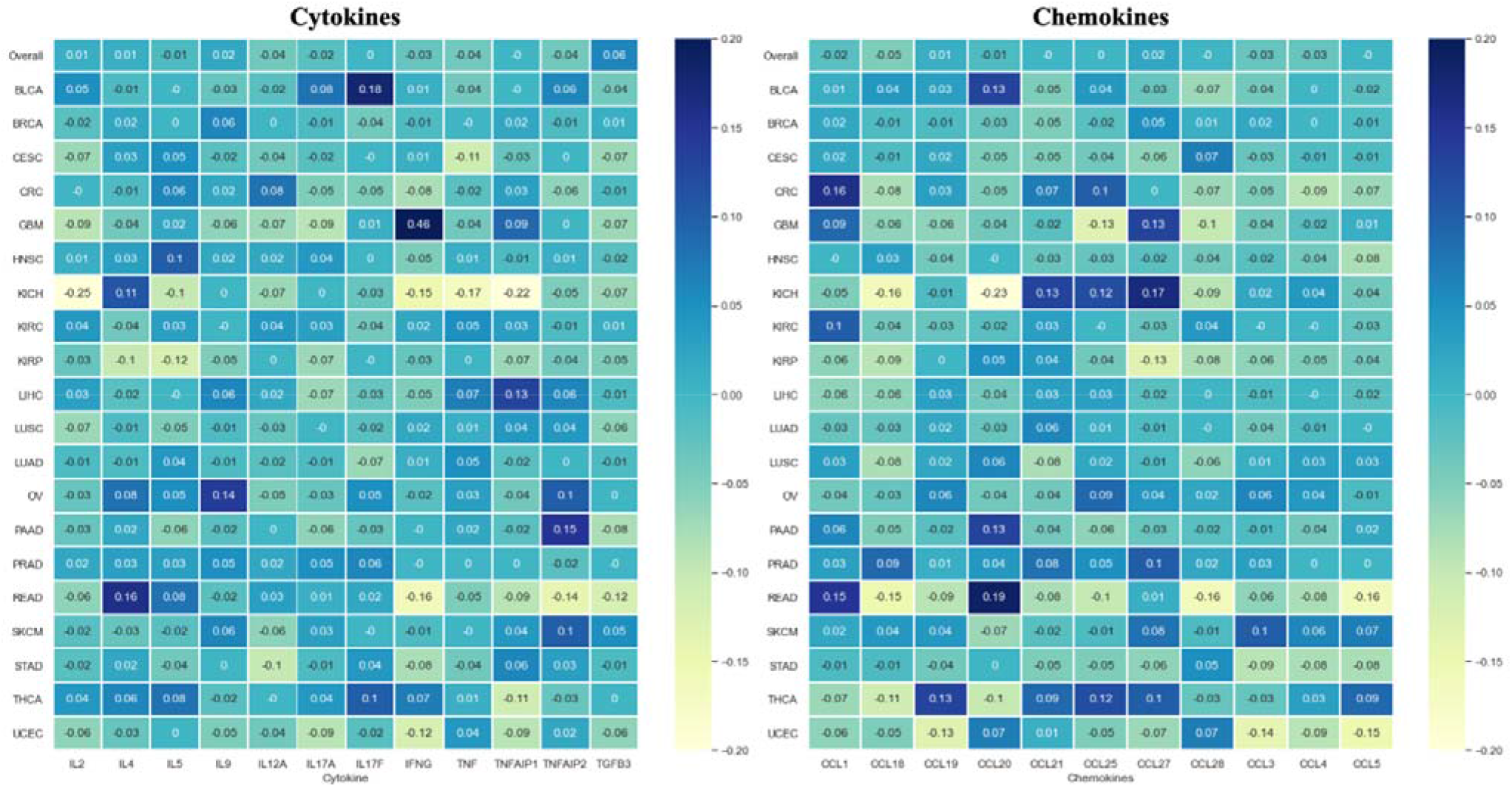
Correlation analysis based on the expression profile of cytokines & chemokines with the overall survival of cancer patients.

### Utility of CancerHLA-I

CancerHLA-I can be interactively browsed and searched in a variety of different ways to satisfy the query of the user. The homepage of CancerHLA-I website provides a simple search page, where users can search query in the database for specific cancer type, HLA-allele, neoantigens, cytokine/chemokine, and its survival association (See Figure 7). The advanced search page in the database provides customized search facility for user defined query using Boolean expressions (AND/OR). We compiled the data in a tabular form corresponding to each cancer type for easy and efficient access. The advantage of the browsing facility is that users can quickly obtain all the results by clicking onto specific entry under concerned category. Moreover, ‘Help’ page on the website provides detailed visualization of the usage of the CancerHLA-I database. The data can be downloaded as a tab-delimited, comma separated, JSON, PDF and XLSX file formats.

**Figure 7:**
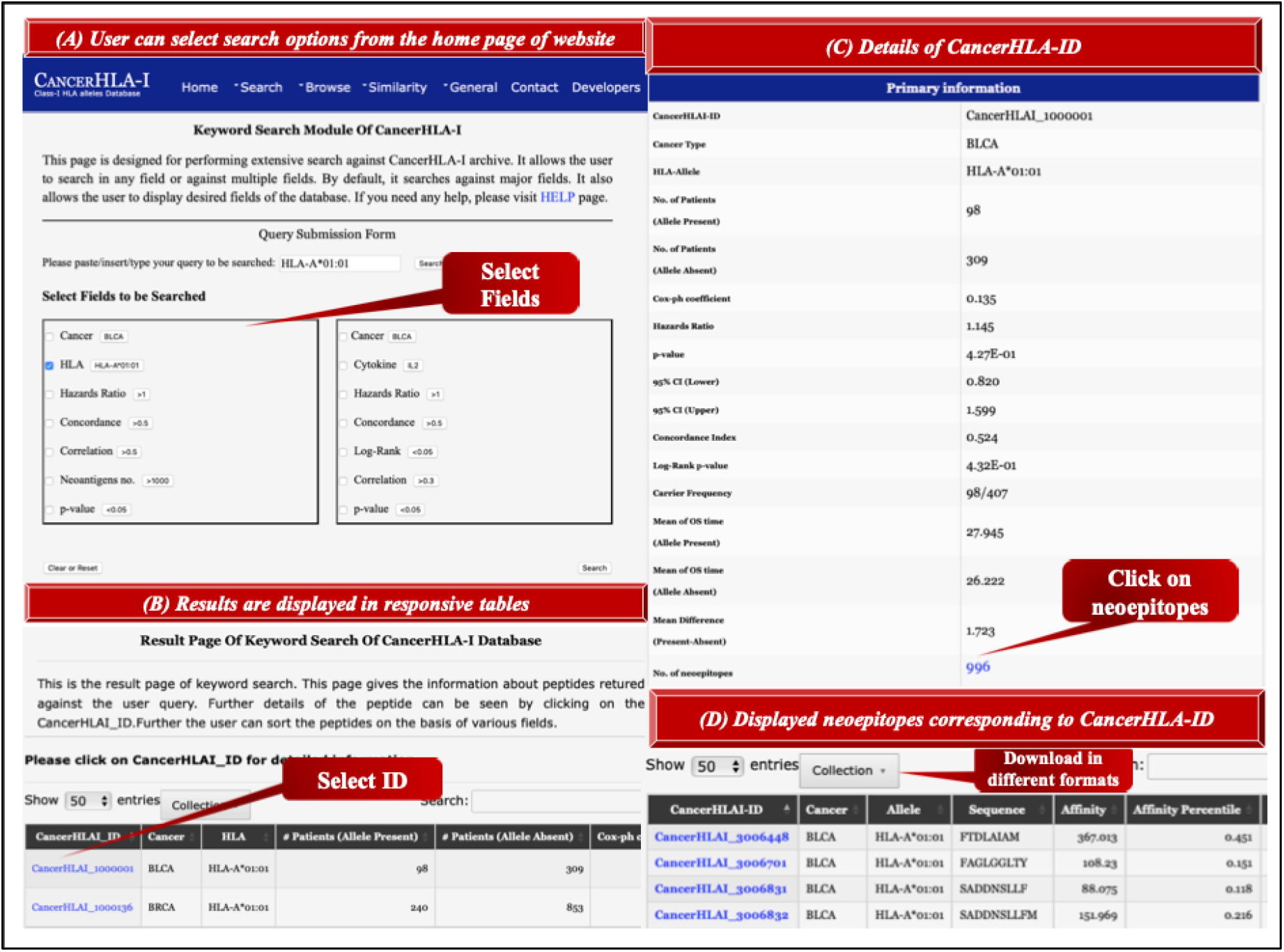
Schematic illustration of the utility of CancerHLA-I (A). Simple search page of the database CancerHLA-I, where users can select fields for querying against the database. Here we have shown the results for the HLA-A*01:01 in the quick search box (B). Complete results against this HLA-alleles are displayed in a responsive table (C). Users can select ID to get detailed information against this query (D). Users can also click on the neoepitopes and download HLA specific binders in the different file formats

## Discussion and Conclusion

Class-I (HLA-A, HLA-B and HLA-C) molecules are essential for immunosurveillance and cancer immunotherapy (Sabbatino et al., 2020;Hazini et al., 2021). It is crucial to present tumor specific peptides or neoantigens via HLA-alleles for the detection and killing of tumor cells by our immune system (van den Bulk et al., 2018;Jiang et al., 2019;Peng et al., 2019;Zhang et al., 2021). However, the loss of the functions of class-I HLA molecules exhibit escape mechanism by different cancer types (Garrido et al., 2016;Dhatchinamoorthy et al., 2021;Hazini et al., 2021). Studies also report that upregulation of class-I non-classical HLA molecules play important role in the cancer immune escape (Bukur et al., 2012;Kochan et al., 2013). Due to mutations at genetic and epigenetic levels loss of heterozygosity occurs in HLA genes at chromosome 6 in non-small cell lung carcinoma and colorectal cancer patients (Hazini et al., 2021;Zhang and Sjoblom, 2021). A recent study reveal that the presence of specific HLA-allele can alter the effect of therapy in different cancers. Researchers observed the presence of HLA-A*03 allele in kidney cancer patients results into reduced survival and poor respond against immune checkpoint blockade (ICI) therapy (Naranbhai et al., 2022). In addition, previous studies report that HLA-B*55 and HLA-A*01 significantly improves the survival while HLA-B*50 allele reduces the survival rate in melanoma patients (Dhall et al., 2020). Moreover, mutations in type-I and II interferon pathway genes also effect the survival of cancer patients. Interleukins such as *IL-6, IL-11, IL-1*, and *TGF-*β induces cancer cell proliferation and progression (Esquivel-Velazquez et al., 2015). Studies reveals that cytokines play important role in the regulation of tumor microenvironment.

Therefore, it is crucial to understand the prognostic role of HLA-alleles and cytokines in order to know the impact of efficacy of cancer immunotherapy. In the present work, we have conducted a study to investigate the connections of Class-I HLA alleles with cancer patient survival in order to aid researchers. We performed pan-cancer analysis on more than 8000 patients in 20 different cancer types. The dataset used in this study obtained from TCGA and TCIA repositories. We used survival data to determine the association between the presence or absence of the 352 unique HLA-alleles. Furthermore, we investigate the relationship between cytokine expression with the overall survival in cancer patients. We observed that the HLA-A*02:01, HLA-A*68:01, HLA-B*52:01, HLA-C*03:02 associated with the poor survival and HLA-C*14:01, HLA-C*12:03 and HLA-A*03:01 improves the survival of cancer patients. Moreover, correlation analysis revealed the positive and negative association of neobinders with the survival rate of the cancer patients. Moreover, we also identified some of the high expression levels of cytokines *IL2, IFNG, IFNB, TNF* significantly improves the survival of cancer patients while some of the cytokines like *IL5, IL17, CCR4, TGF* reduces the survival rate significantly. We incorporated overall results in the highly interactive web-based platform for the analysis and identification of cancer-specific biomarkers. We have integrated user-friendly browsing, searching, and analysis modules in our resource “CancerHLA-I” (https://webs.iiitd.edu.in/raghava/cancerhla1/) for easy data retrieval, data comparison, and examination. We anticipate that our research yields promising novel HLA and cytokines based biomarkers for improved cancer immunotherapy and treatment.

## List of Abbreviations

BLCA: bladder urothelial carcinoma
BRCA: beast invasive carcinoma
CESC: cervical squamous cell carcinoma and endocervical adenocarcinoma
CHOL: cholangiocarcinoma
GBM: glioblastoma multiforme
HNSC: head and neck squamous cell carcinoma
KICH: kidney chromophobe
KIRC: kidney renal clear cell carcinoma
KIRP: kidney renal papillary cell carcinoma
LIHC: liver hepatocellular carcinoma
LUAD: lung adenocarcinoma
LUSC: lung squamous cell carcinoma
OV: ovarian serous cystadenocarcinoma
PAAD: pancreatic adenocarcinoma
PRAD: prostate adenocarcinoma
READ: rectum adenocarcinoma
SKCM: skin cutaneous melanoma
STAD: stomach adenocarcinoma
THCA: thyroid carcinoma
UCEC: uterine corpus endometrial carcinoma

## Authors’ contributions

AD and GPSR collected and processed the datasets. AD, HK, SP and GPSR conceived the idea and analysed the results. AD and SP created the back-end of the web server the front-end user interface. AD, SP and GPSR penned the manuscript. GPSR conceived and coordinated the project. All authors have read and approved the final manuscript.

## Acknowledgements

Authors are thankful to the Department of Bio-Technology (DBT) and Department of Science and Technology (DST-INSPIRE) for fellowships and the financial support and Department of Computational Biology, IIITD New Delhi for infrastructure and facilities.

